# TCRAD: An End-to-End Framework for Antigen-Targeted T Cell Receptor Design

**DOI:** 10.64898/2026.01.21.700513

**Authors:** Chenao Li, Yaochi Guo, Xin Guan, Hui Chen, Yong Zhang, Pengyuan Yang, Jizhong Lou

**Affiliations:** State Key Laboratory of Epigenetic Regulation and Intervention, CAS Center for Excellence in Biomacromolecules, Institute of Biophysics, Chinese Academy of Sciences, Beijing 100101, China; University of Chinese Academy of Sciences, Beijing 100049, China

## Abstract

Understanding and engineering T-cell receptor (TCR) specificity is central to personalized immunotherapy and antigen discovery. However, while antigen-conditioned TCR design approaches have begun to emerge, most frameworks still focus on scoring candidate TCR–pMHC pairs, and achieving high hit rates in designing functional, target-reactive TCRs remains a key bottleneck. Here, we present TCRAD, a deep learning pipeline that performs de novo design of TCR CDR3β sequences conditioned on antigenic peptides. TCRAD comprises three modules: Sequence Generation, Sequence Filtration, and Structure Prediction. TCRAD can effectively generate antigen-specific CDR3β sequences, discriminate functional binders with high accuracy, and predict bound and unbound structures with performance comparable to or exceeding state-of-art methods. Experimental validation using the 1G4/NY-ESO-1 system demonstrated that the model can design novel functional TCRs capable of surface expression, specific pMHC binding, and that 17.2% candidates (5 of 29) elicit antigen-induced activation. Together, these results establish TCRAD as a powerful framework for TCR sequence design, offering a scalable path toward rational engineering of antigen-specific TCRs for cancer immunotherapy, vaccine development, and personalized immunotherapeutics.

## Introduction

The T cell receptor (TCR) is central to adaptive immunity and underlies the specificity of T-cell responses in infection, cancer, and autoimmunity^1, 2, 3, 4^. TCRs recognize antigenic peptides presented by major histocompatibility complex (MHC) molecules, initiating downstream signaling that determines T-cell activation, differentiation and effector function ^5,6^. The diversity of TCR recognition stems from the combinatorial assembly of variable, diversity, and joining (V(D)J) segments and extensive junctional variability, producing millions of unique TCRαβ or TCRγδ sequences capable of responding to a broad antigenic landscape^7, 8^.

Among the complementarity-determining regions (CDRs) and framework regions (FRs) of TCRs, the CDR3β (CDR3 on the β chain) is the most diverse and plays a dominant role in contacting the peptide within the pMHC complex ^9, 10^, a detailed understanding of its sequence-structure-function relationships is therefore crucial for rational TCR design and for developing personalized immunotherapies such as TCR-engineered T-cell (TCR-T) therapy^11^.

Recent years have seen the rapid development of deep learning models for predicting TCR-epitope or pMHC binding. Methods such as Pan-Peptide^12^, TEIM^13^, pMTattn^14^, TransPHLA^15^, TCR-BERT^16^, SC-AIR-BERT^17^, and DeepAIR^18^ have introduced a variety of architectures from meta-learning^19, 20, 21^, neural Turing machines^22^ to cross-attention and Transformer^23^ to improve binding prediction accuracy. These models are discriminative predictors that score candidate TCR–peptide/HLA pairs.

More recently, antigen-conditioned generative approaches for TCR design have begun to appear^24, 25^; however, end-to-end workflows that couple generation with rigorous downstream filtration remain limited. Consistent with this gap, prior design studies have reported low hit rates (1/40, 2.5%) in functional assays^24^, underscoring the need for more reliable methods with higher empirical success rates. With the emergence of powerful generative AI frameworks ^26, 27, 28, 29^ and their widespread application in biomedical field^30, 31, 32^, there is growing opportunity to develop automated pipelines for TCR sequence design.

In this study, we present TCRAD, an automated deep-learning pipeline that generates, filters, and evaluates CDR3β sequences based solely on antigenic peptide input. TCRAD integrates (i) a UniLM-based generation model capable of flexible mask-prediction strategies; (ii) two BERT-based^33^ discriminators that assess natural plausibility and antigen specificity; and (iii) a structure prediction module inspired by Helixfold and ESM-fold^34, 35^ that operates without multiple sequence alignments. We benchmarked each module against state-of-the-art methods and validated the full pipeline using biological experiments. TCRAD achieves superior binding discrimination in zero-shot evaluations, generates structurally plausible CDR3β conformations, and enables the de novo design of functional TCRs against the NY-ESO-1 antigen using the 1G4 framework. Our findings highlight the ability of TCRAD to accelerate TCR engineering and provide a foundation for next-generation personalized immunotherapies.

## Results

### TCRAD Enables Antigen-Specifically Generation of CDR3β

TCRAD consists of three coordinated modules: Sequence Generation (SG), Sequence Filtration (SF) and Structure Prediction (SP) (Fig. 1). The SG Module is built upon UniLM^26^, a unified language model capable of flexible mask-based prediction. The UniLM model demonstrates versatility in handling various tasks related to sequence generation and semantic understanding^26^. By manipulating its masked attention patterns, UniLM can perform a variety of generative and predictive tasks. In our pipeline, UniLM receives an antigenic peptide sequence as input and iteratively generate corresponding CDR3β in a predefined positional order (Fig. 2).

**Fig. 1:**
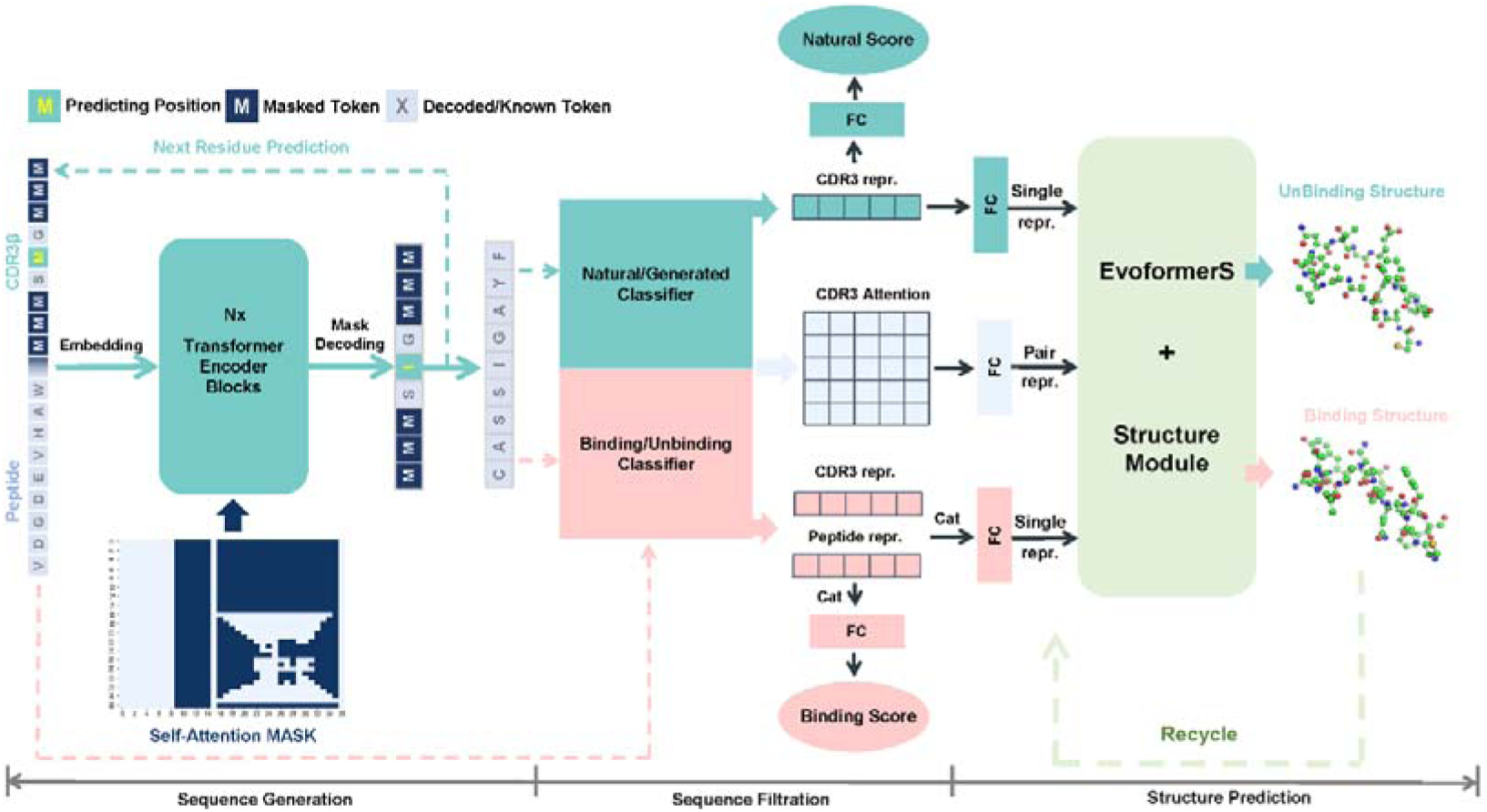
Overview of the TCRAD framework for antigen-specific TCR CDR3β design. Schematic overview of the TCRAD pipeline comprising three coordinated modules: Sequence Generation (SG), Sequence Filtration (SF), and Structure Prediction (SP). Given a peptide antigen, the SG module generates candidate CDR3β sequences using multiple attention-mask–based generation strategies. The SF module evaluates generated sequences using binding and naturalness classifiers to retain high-confidence candidates. The SP module subsequently assesses structural compatibility by modeling the local CDR3β loop structure of the filtered sequences.

**Fig. 2:**
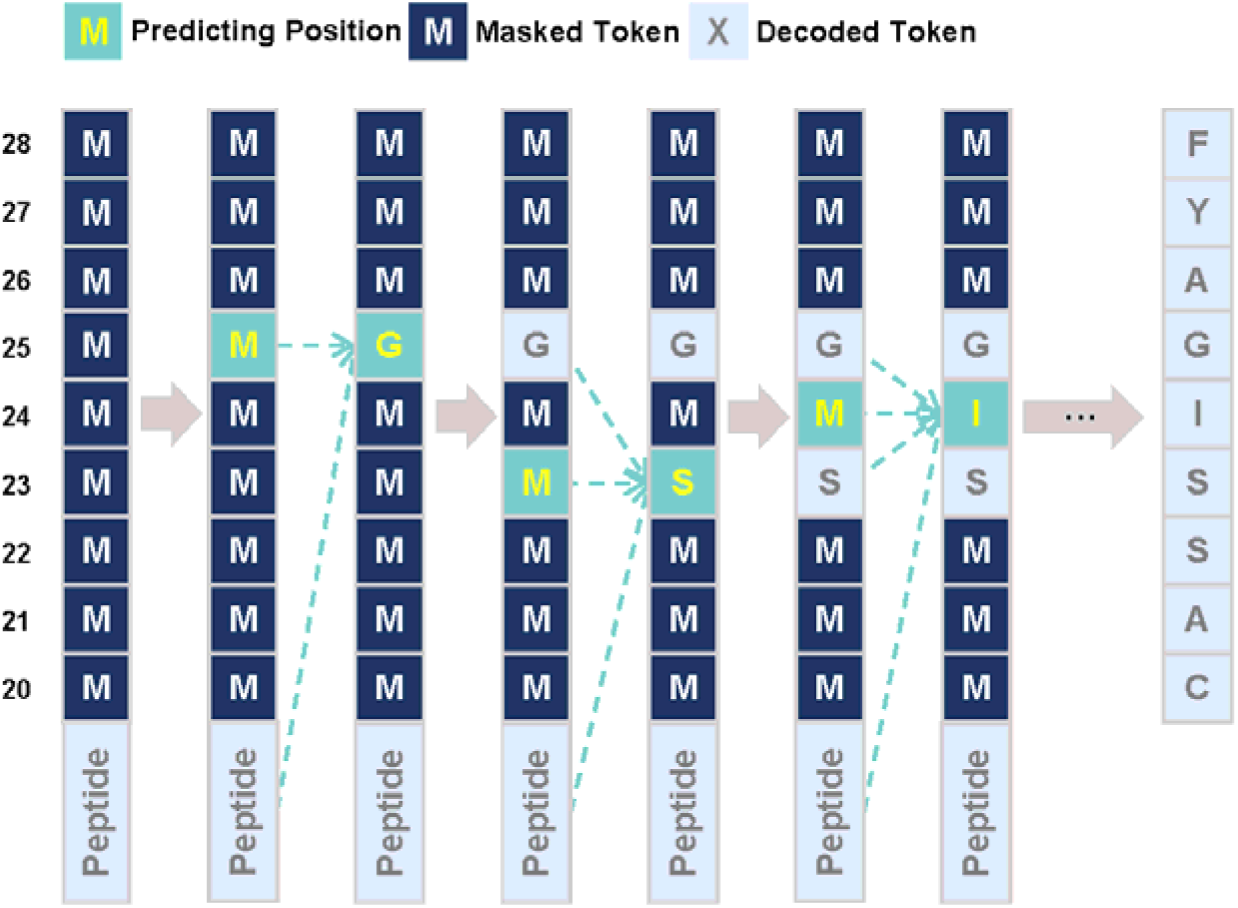
Process of Generating a CDR3β sequence. The model starts from a fully masked CDR3β segment and iteratively selects one residue position to unmask and predict, conditioned on the peptide and previously decoded CDR3β residues. At each step, the target position attends to the peptide and past CDR3β tokens but not to future positions, enforcing a causal masked self-attention pattern until the complete CDR3β sequence is generated.

Given the limited prior evidence on the optimal peptide generation direction, and because central CDR3β residues often play outsized roles in antigen recognition^4, 36, 37, 38^, we implemented six sequence generation orders, each defined by a distinct attention-masking strategy (Fig. 3). Rather than selecting a single generation order, we retain multiple complementary strategies to improve robustness and diversity, as different CDR3β residues can dominate peptide recognition across antigen contexts.

**Fig. 3:**
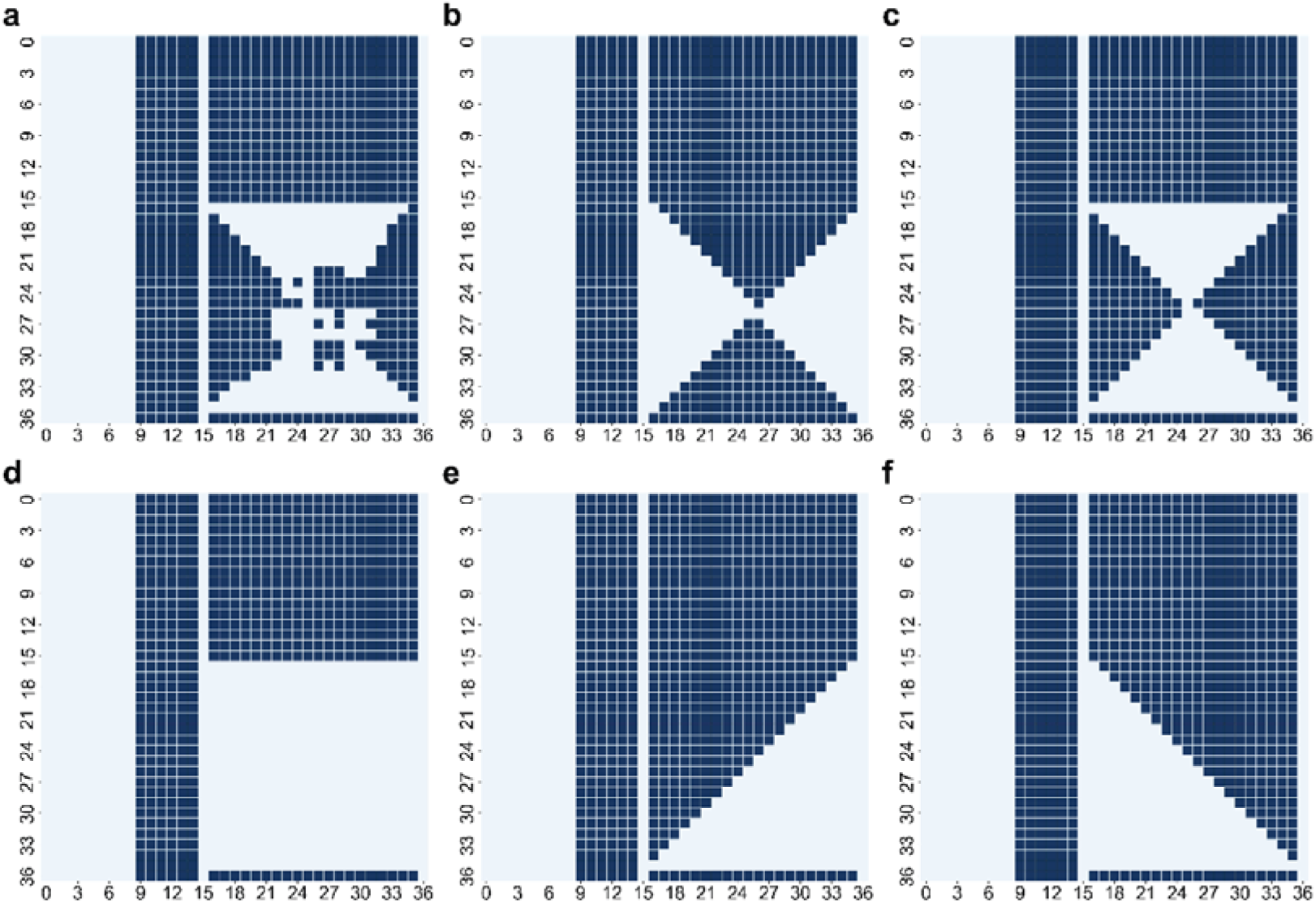
Masking strategies for CDR3β sequence generation. **a.** Left-to-Right (L2R) generation. **b.** Right-to-Left (R2L) generation. **c.** Bidirectional (Bidirection) generation with full contextual information. **d.** End-to-center (RL2M) generation from both termini toward the central region. **e.** Center-outward (Middle) generation from the central region toward both termini. **f.** Priority-based (Prioritization) generation following a predefined importance ranking of CDR3β positions.

To enhance the multi-task adaptability of the CDR3β-generation model, we trained the model on a dataset comprising 157,000 peptide-CDR3β pairs with experimentally verified positive reactivity (K1-positive, Methods). During training, the model learned six different mask prediction tasks on this training set, enabling it to infer masked amino acids based on the peptide and various observed CDR3β contexts.

The performance for each generation orders is summarized in Table 1, which reports the top-k accuracies for masked-residue prediction. Table 2 provides representative examples of designed CDR3β sequences targeting the NY-ESO-1 antigen in the 1G4 TCR framework.

**Table 1.**
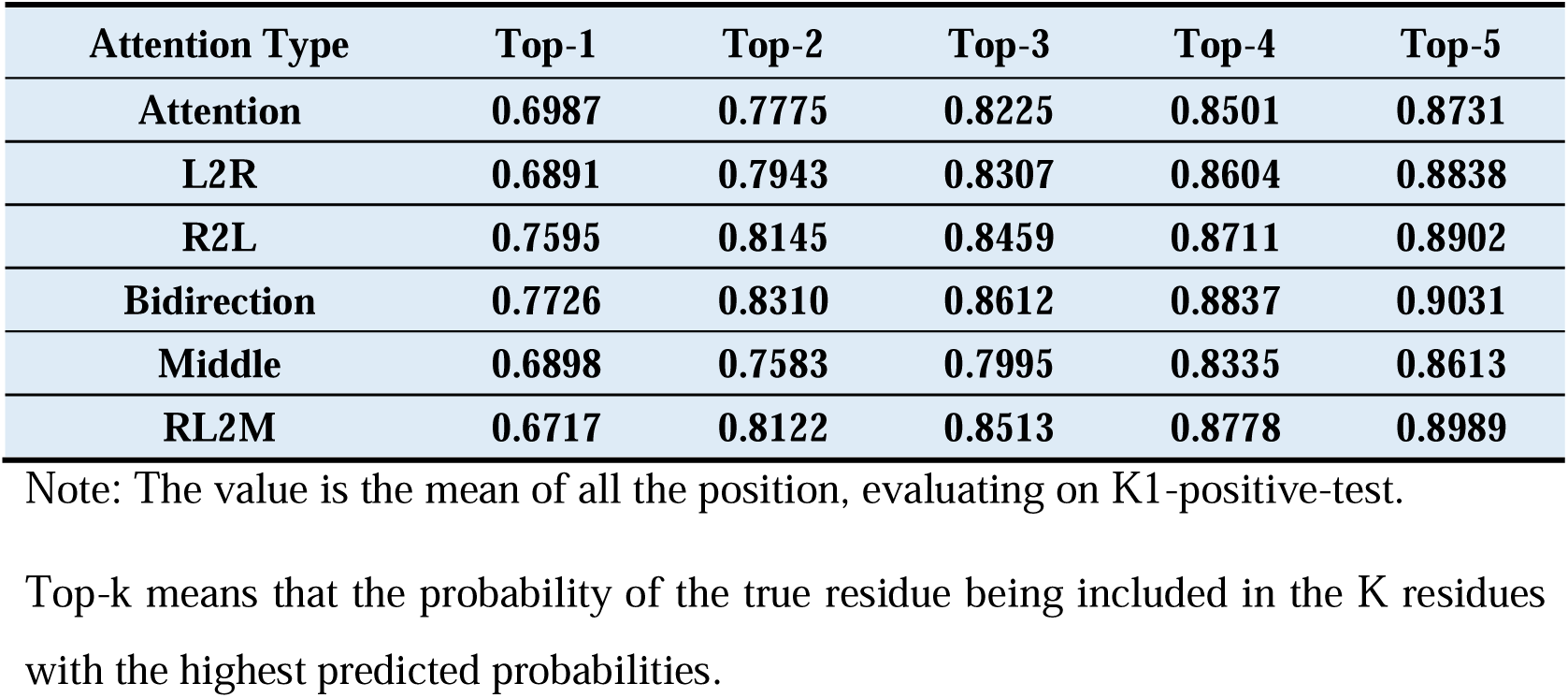
The Top-k Accuracy of the Sequence Generation Module.

| Attention Type | Top-1 | Top-2 | Top-3 | Top-4 | Top-5 |
| --- | --- | --- | --- | --- | --- |
| Attention | 0.6987 | 0.7775 | 0.8225 | 0.8501 | 0.8731 |
| L2R | 0.6891 | 0.7943 | 0.8307 | 0.8604 | 0.8838 |
| R2L | 0.7595 | 0.8145 | 0.8459 | 0.8711 | 0.8902 |
| Bidirection | 0.7726 | 0.8310 | 0.8612 | 0.8837 | 0.9031 |
| Middle | 0.6898 | 0.7583 | 0.7995 | 0.8335 | 0.8613 |
| RL2M | 0.6717 | 0.8122 | 0.8513 | 0.8778 | 0.8989 |

Note: The value is the mean of all the position, evaluating on K1-positive-test.
Top-k means that the probability of the true residue being included in the K residues with the highest predicted probabilities.

**Table 2:** CDR3β Redesign and Mutation based on NY-ESO-1 antigen.

| Input Peptide | Design Mode | Input CDR3β | Generated CDR3β | Binding Score (>0.28) | Natural Score (<0.56) |
| --- | --- | --- | --- | --- | --- |
| SLLMWITQC | Full Design | ----- | CASSGGTSTNYNEQFF | 0.9158 | 0.1972 |
|  |  |  | CSALGLGAGGSGNTIYF | 0.7758 | 0.1399 |
|  | Mutation Optimization | CASSY----GELFF | CASSYRAGGGELFF | 0.8285 | 5.72E-05 |
|  |  |  | CASSYRRGGGELFF | 0.7043 | 2.35E-05 |

Note: "-" represents the positions to be generated.
Full Design refers to designing the entire CDR3β sequence across all positions. Mutation Optimization indicates mutating selected positions of the CDR3β . Binding Score and Natural Score range from 0 to 1: higher Binding Score indicates higher binding likelihood, whereas lower Natural Score indicates greater CDR3β naturalness.

### Dual Filtration Ensures Antigen Specificity and Naturalness of Generated CDR3β

Following sequence generation, TCRAD applies the SF module to assess the functional plausibility of each generated CD3β candidate. We employ two discriminators to assess the validity of the candidate sequences: reaction specificity and biophysical probability. Reaction specificity pertains to whether the generated CDR3β can bind specifically to the antigen. Biophysical probability refers to the generated CDR3β sequences being physically plausible in their natural state, adhering to the physical laws of the objective world. We simplify this problem to evaluating the disparity between the generated CDR3β sequences and those found in nature.

Thus, the SF module consists of two Bert-based classifiers: Binding Classifier and Naturalness Classifier. The binding classifier evaluates whether a generated CDR3β is likely to bind the input peptide, it takes a peptide-CDR3β pair as input, outputting feature representations for the pair. These representations are then fed into a Multi-Layer Perceptron (MLP) to obtain a binding score. The naturalness classifier assesses whether a generated CDR3β resembles naturally occurring TCR sequences, it only takes a CDR3β sequence for feature extraction, and the extracted features are input into an MLP to obtain a naturalness score. Both models generate self-attention weight matrices that capture the residue-level dependencies and serve as geometric features for downstream structural modeling.

To train these models, we assembled three datasets, K1, K2, and K3. Dataset K1 comprising 519 k instances of peptide-CDR3β binding data. Dataset K2 comprising 12 M instances of natural/generated CDR3β sequences. Dataset K3 is an independent peptide–CDR3β dataset for zero-shot benchmarking (Supporting Table S1).

The Binding Classfier performed strongly on both the K1 training and K1 test sets, as measured by AUROC and AUPR (Fig. 4a). The classification threshold of both classifiers was determined using the ROC curve (Fig. 4b). Zero-shot evaluation on K3 dataset showed that our binding classifier outperformed state-of-the-art TCR-pMHC binding prediction models including TEIM^13^, TABR-Bert^39^, Panpep^12^, pMTnet^40^. The results, illustrated in ROC and PR curves (Fig. 4c,d), demonstrated a significant advantage of our model. See supplemental methods for the details of the implementation of these methods.

**Fig. 4:**
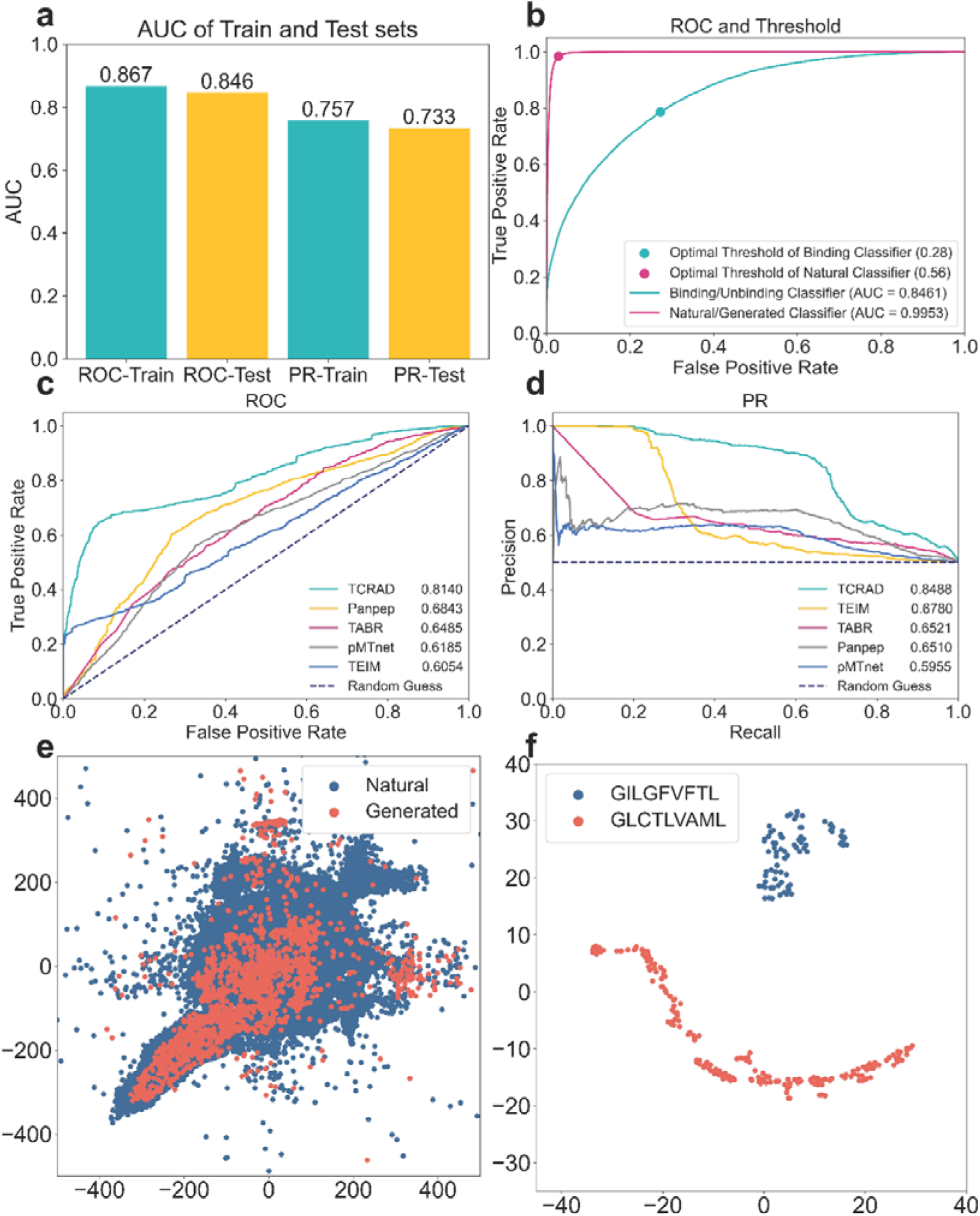
Performance of the Sequence Filtration Module. **a.** AUROC and AUPR curves of the Binding Classifier evaluated on the K1-train and K1-test sets. **b.** ROC curves and selected decision thresholds for the Binding Classifier and Naturalness Classifier, evaluated on the K1-test set and the K2-test set. **c**, **d**. ROC (c) and PR (d) curves comparison of the Binding Classifier and SOTA baseline methods on the K3 data set. **e**. Feature-space visualization of natural CDR3β sequences and the CDR3β sequences generated by SG module, evaluated by the Naturalness Classifier using datasets of 1 million generated and 5.5 million natural sequences. **f**. Visualization analysis for CDR3β sequences designed for the epitopes GLCTLVAML and GILGFVFTL.

To evaluate whether the SG module produces biologically realistic CDR3β sequences, we projected both natural and generated CDR3β into the feature space learned by the Naturalness Classifier and visualized them using T-SNE^41^. We constructed a large-scale reference set by augmenting the K2 dataset with an additional 4.5 million collected natural CDR3β sequences. In this embedding space, the generated CDR3β were largely confined within the broad cloud of natural repertoires (Fig. 4e), indicating that the SG Module inherently produces CDR3β that resemble natural patterns.

Additionally, we performed design for two Peptide-CD3β pairs, GILGFVFTL-CASSTRAGDTQYF and GLCTLVAML-CASSQSPGGTQYF (sourced from NetTCR-2.0^42^). Specifically, we retained the starting CASS and ending QYF residues of the CDR3β sequence, allowing the SG module to design the amino acid residues located in the middle. The results revealed that the generated sequences cluster distinctly by peptide (Fig. 4f, and Supporting Tables S2 and S3), confirming that TCRAD can design peptide-specific CDR3β variants.

Based on the trained Binding Classifier, we performed an analysis of the positional importance during the peptide-CDR3β binding prediction on the test set. The attention matrix of the peptide-CDR3β (Fig. 5a) was extracted and the positional weights of CDR3β (Fig. 5b) was subsequently summed and normalized. The attention weights distribution for CDR3β exhibited a trapezoidal shape, with central residues contributing the most to peptide recognition. This observation further supports the design of multi-order generation strategies in the SG module. Quantitative analysis of positional importance allowed us to derive a CDR3β sequence generation order based on importance, where more crucial positions were generated earlier.

**Fig. 5:**
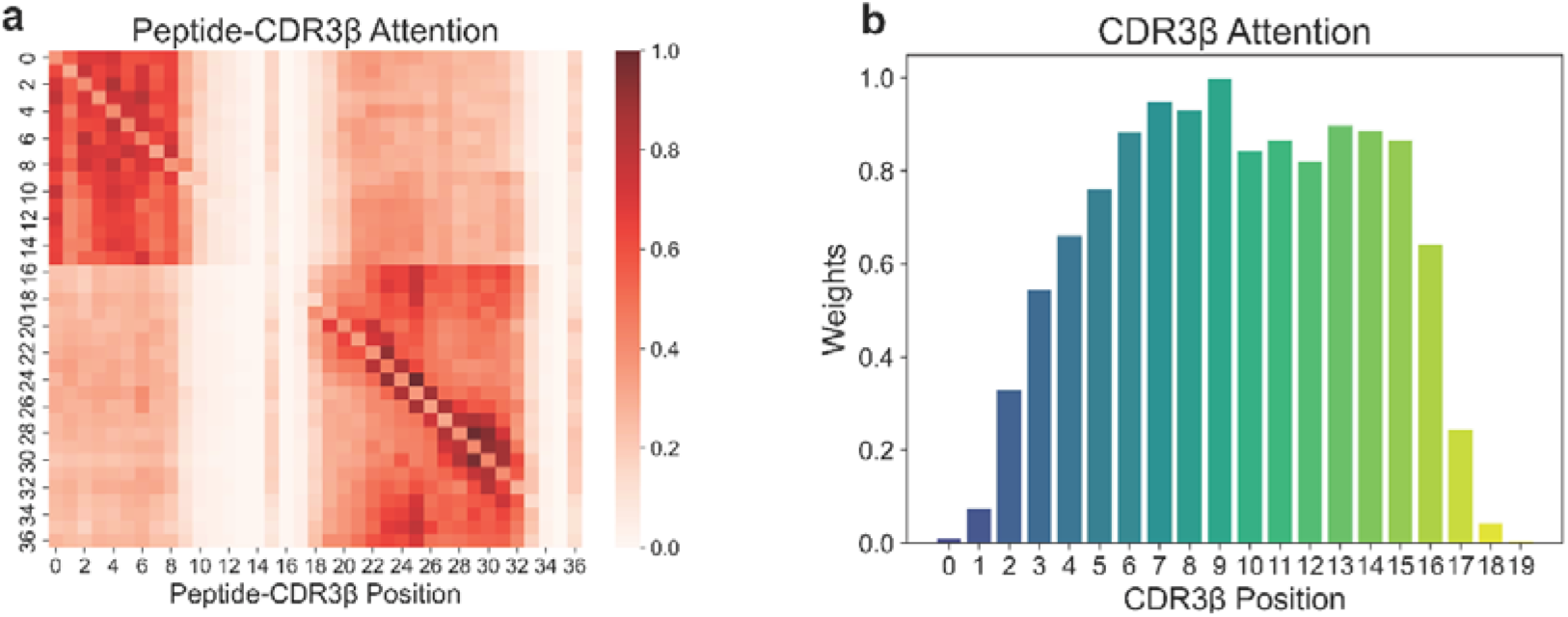
TCR–epitope binding analysis. **a.** Attention weight distribution between peptide residues and CDR3β residues in the peptide–CDR3β binding task. **b.** Positional distribution of attention weights across CDR3β residues, highlighting relative contribution of individual positions to peptide recognition.

Together, these results demonstrate that the SF module effectively filters raw outputs from the SG module and ensures that downstream structural modeling operates on realistic and antigen-relevant candidates.

### Structure Prediction Module Accurately Models CDR3β Conformations in Bound and Unbound States

After filtration, selected CDR3β sequences are passed to the SP module, which estimates their three-dimensional conformations. The SP module was inspired primarily from the work Helixfold and ESM-fold ^34, 35^, where a language model is used in place of multiple sequence alignment (MSA) part. Specifically, sequence embeddings extracted by the language model are fed into the Evoformer blocks to model residue–residue interactions, and the resulting representations are then decoded by a structure module to produce atomic coordinates for the predicted complex.

This SP module supports two inference modes: the Unbound mode and the Bound mode. The Unbound mode uses CDR3β features derived from the Naturalness Classifier to predict the unbinding conformation of the CDR3β region and the Bound mode incorporates peptide–CDR3β joint features from the Binding Classifier for the prediction of the binding conformation of CDR3β region with the peptide. These complementary modes allow TCRAD to model CDR3β both in isolation and within its peptide-engaged conformation.

We benchmarked the SP module against TCRmodel2^43^, which based on Alphafold-Multimer^44^ model. Across the test set, TCRAD achieved an average RMSD of around 1 Å for both bound and unbound predictions (Fig. 6a,b). Notably, the SP module consistently outperformed TCRmodel2 in the bound state, highlighting the advantage of integrating peptide-aware features extracted by the Binding Classifier.

**Fig. 6:**
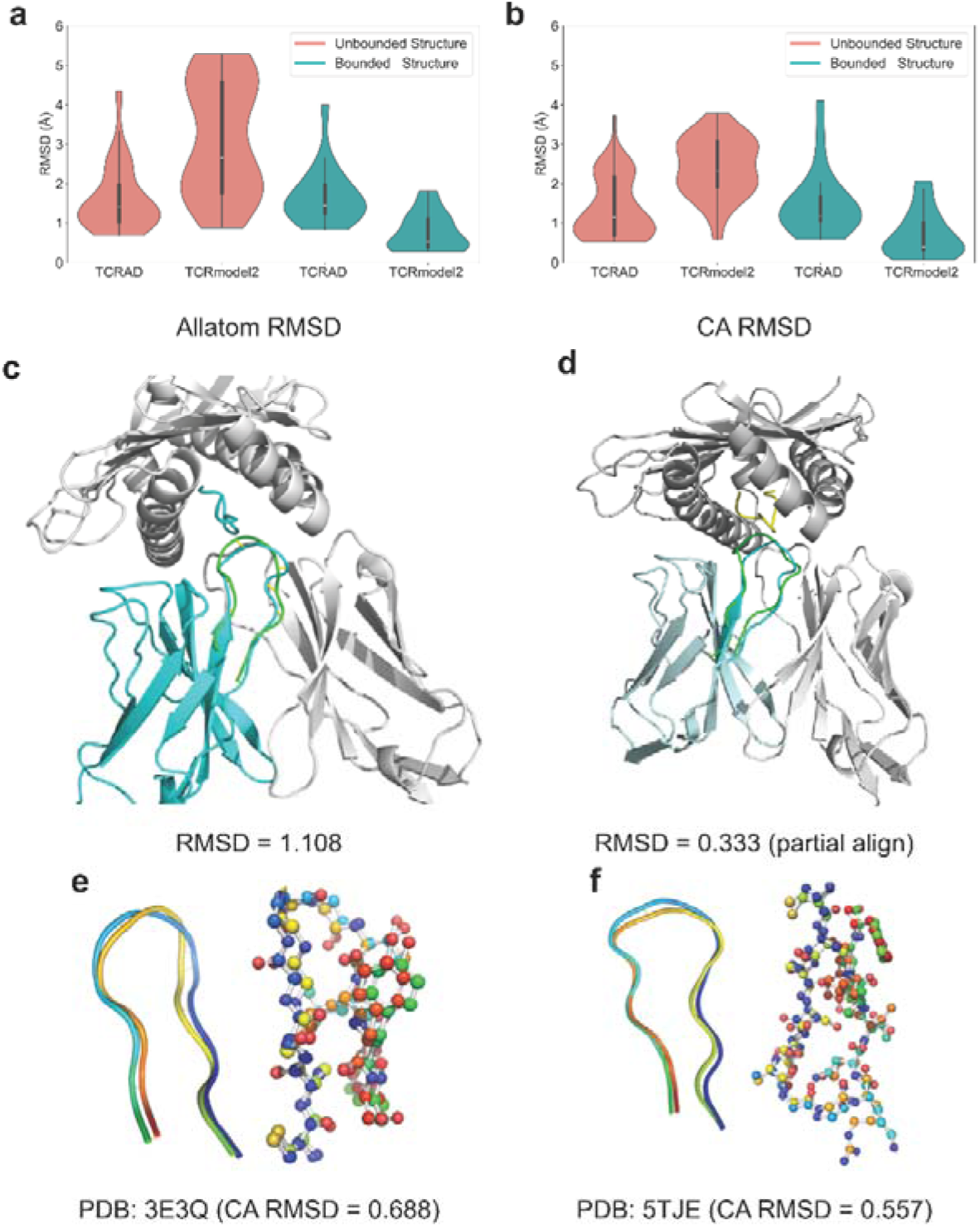
Comparison of CDR3β structure prediction with TCRmodel2. **a**, **b**. Root-mean-square deviation (RMSD) calculated over all atoms (a) or Cα atom only (b) for predicted CDR3β structures. From left to right, columns show: bound structures predicted by the SP Module, bound structures predicted by TCRmodel2, unbound structures predicted by the SP Module, and unbound structures predicted by TCRmodel2. **c.** Predicted CDR3β structure (green) for PDB entry 2BNR (sequence: CASSYVGNTGELFF) aligned with the experimentally resolved structure (cyan). **d.** Predicted structure (green) of a designed CDR3β sequence (CASSGGTSTNYNEQFF) for PDB entry 2BNR in complex with the peptide SLLMWITQC (cyan), aligned using the same N-terminal Cα and C-terminal phenylalanine residues. **e.** Representative CDR3β structure prediction for PDB entry 3E3Q. **f.** Representative CDR3β structure prediction for PDB entry 5TJE.

These results demonstrate that the SP module provides accurate and context-sensitive structural predictions, forming a strong foundation for evaluating designed sequences prior to experimental validation.

### Experimental validation of De Novo Designed CDR3β sequences 1G4/NY-ESO-1 System

To experimentally validate TCRAD, we focused on the clinically relevant NY-ESO-1 / HLA-A2 antigen pair and the canonical 1G4 TCR. The NY-ESO-1 protein was first identified as a tumor-associated antigen aberrantly expressed in cancer cells, through an autologous antibody screening method,^45^ it belongs to the cancer-testis antigen (CT antigen) family and is one of the most immunogenic tumor antigens discovered to date with broad tumor expression^46^. Although the 1G4 TCR selectively recognizes the HLA-A2/NY-ESO-1_157-165_ (SLLMWITQC) peptide complex, making it valuable target for the development of NY-ESO-1-based immunotherapy^47, 48, 49, 50^, affinity-enhanced variants have demonstrated off-target liabilities^51, 52, 53, 54, 55, 56^, underscoring the need for safer, sequence-diverse alternatives.

We used TCRAD to design 29 unique CDR3β sequences targeting HLA-A2/NY-ESO-1, and grafted them into the 1G4 TCR framework (Table S5). Lentiviral transduction introduced these TCRs into J76 cells lacking endogenous TCR expression. Flow cytometry revealed that 8 designed TCRs displayed robust surface expression and bound efficiently to NY-ESO-1/HLA-A2 tetramers (Fig. 7a,b). Notably, these 8 tetramer-positive designs extend beyond NY-ESO-1 –associated CDR3β sequences in the training set K1-positive(minimum Levenshtein distance: 3; mean: 5.125).

**Fig. 7:**
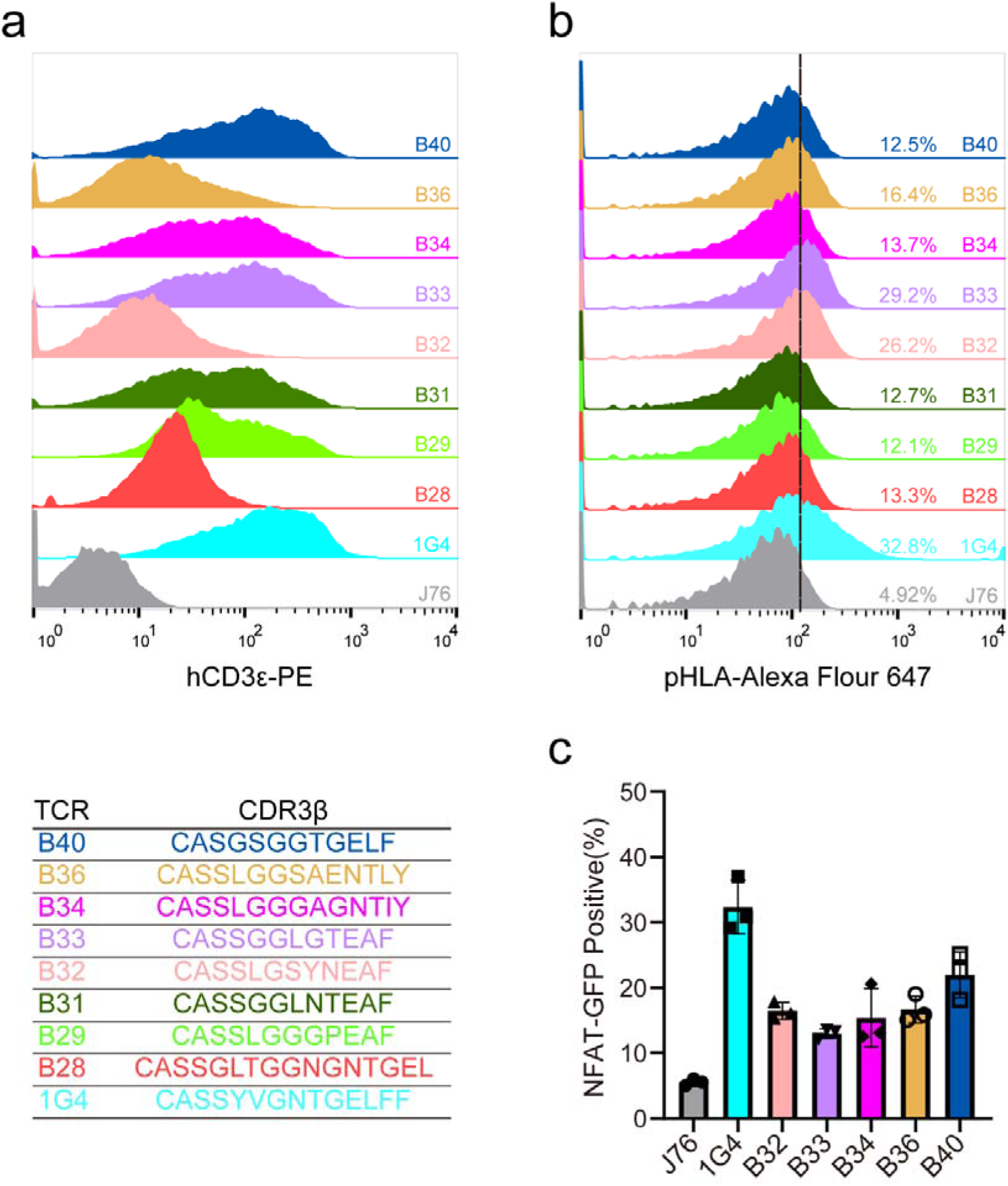
Experimental validation of designed TCRs. **a.** Flow cytometry analysis of TCR surface expression in J76 cells expressing either wild-type (1G4) or designed TCRs. **b.** The binding of designed TCRs to NY-ESO-1/HLA-A*02:01 (pHLA) complexes. **c.** NFAT–GFP reporter assay quantifying functional TCR activation in response to pHLA stimulation.

To assess functionality, the engineered TCRs were expressed in NFAT-reporter J76 cells and stimulated with NY-ESO-1/HLA-A2. Five designed TCRs mediated strong NFAT activation upon peptide exposure (Fig. 7c), confirming that sequences generated by TCRAD are capable of forming functional antigen-specific TCR complexes.

These results provide direct biological evidence that TCRAD can design diverse, structurally plausible, and functional CDR3β sequences that retain specificity while expanding sequence space beyond naturally observed repertoires.

## Discussion

In this study, we introduce TCRAD, a unified deep-learning pipeline capable of de novo generation, filtration, and structural evaluation of TCR CDR3β conditioned solely on the antigenic peptide. By integrating a UniLM-based generative model with BERT-based discriminators and a peptide-aware structural prediction framework, TCRAD represents a substantial advance beyond previous TCR analysis tools, which have predominantly focused on predicting binding for known TCR-peptide pairs rather than generating new TCRs.

A key contribution of TCRAD is its ability to generate antigen-specific CDR3β sequences in a directionally flexible manner. Unlike traditional left-to-right or fully autoregressive approaches, TCRAD employs six complementary generation patterns, enabling the model to capture position-specific constraints and long-range dependencies that are intrinsic to CDR3β structure and function. This strategy is supported by our attention analyses and downstream filtration results, which highlight the functional importance of central CDR3β residues for peptide recognition. By jointly training six mask-prediction tasks, the Sequence Generation module gains robustness and flexibility, allowing it to produce sequences that are both diverse and tailored to the input antigen.

The SF module further ensures functional plausibility by separately evaluating naturalness and antigen specificity. The strong performance of the Binding Classifier in both standard and zero-shot settings underscores the strength of its learned representation of TCR–peptide interactions. Remarkably, the model surpasses state-of-the-art predictors such as TEIM, TABR-BERT, Pan-Peptide, and pMTnet in zero-shot evaluation, suggesting that the joint embedding of peptide and CDR3β provides a more expressive and generalizable representation of binding determinants. Meanwhile, the Naturalness Classifier anchors the generator to realistic sequence space, as demonstrated by t-SNE analyses showing broad overlap between generated and natural CDR3β distributions. Together, these discriminators create a principled mechanism for refining raw generative output into biologically credible candidates.

Building upon these filtered sequences, the SP module extends TCRAD into the structural domain. Leveraging language-model–derived attention features instead of MSAs, the SP module provides efficient and accurate estimation of both bound and unbound CDR3β conformations. The model outperforms AlphaFold-Multimer–based TCRmodel2 in predicting bound-state configurations, demonstrating that peptide-conditioned embeddings from the Binding Classifier capture local structural determinants critical for TCR–pMHC engagement. This highlights the utility of deeply integrating generative, discriminative, and structural components within a single pipeline, enabling TCRAD to assess candidates through sequence-level, functional, and geometric criteria.

Most importantly, our experimental validation in the 1G4 TCR/NY-ESO-1 system provides direct evidence that TCRAD is not only computationally effective but also biologically meaningful. Several designed CDR3β produced functional TCRs capable of surface expression, pMHC tetramer binding, and 17.2% candidates (5 of 29) further elicited antigen-induced NFAT activation. These results confirm that TCRAD can navigate the vast TCR sequence space and identify viable, antigen-reactive TCR candidates. Considering the known off-target risks associated with affinity-enhanced 1G4 variants, the ability to design structurally plausible and functionally competent CDR3β alternatives underscores the utility of TCRAD for therapeutic development.

Despite these advances, several limitations remain. First, the current pipeline focuses exclusively on CDR3β, omitting contributions from the α-chain and from CDR1/2 loops, which also influence TCR specificity and stability. Expanding TCRAD to generate full TCRαβ sequences—including paired design—represents an essential next step. Second, while our structural model effectively predicts isolated CDR3β conformations, future versions could incorporate energetic scoring functions, molecular dynamics–based refinement, or pMHC docking models to more accurately assess conformational stability and peptide engagement. Third, although our experimental validation demonstrates clear functionality, large-scale screening across diverse epitopes will be necessary to fully assess generalizability and clinical potential.

Looking forward, TCRAD offers exciting avenues for personalized immunotherapy, vaccine design, and immune repertoire engineering. The ability to generate antigen-specific TCRs on demand could accelerate the identification of therapeutic TCRs for cancer neoantigens, viral epitopes, or autoimmune targets. Moreover, by enabling systematic exploration of the TCR sequence–structure–function landscape, TCRAD may help elucidate fundamental principles of TCR recognition that remain elusive due to the enormous combinatorial diversity of TCR repertoires.

In summary, TCRAD represents a significant step toward fully automated TCR design, moving beyond predictive modeling to enable controlled, antigen-conditioned generation and evaluation of novel TCR candidates with a high experimental hit rate. As deep generative models continue to advance, we anticipate that pipelines like TCRAD will play an increasingly central role in the rational engineering of adaptive immune receptors and the development of next-generation precision immunotherapies.

## Data Availability

The datasets can be accessed on Zenodo (https://zenodo.org/records/10715856). This includes the sequence-based datasets mentioned in this paper, namely K1, K2, and K3 datasets. Additionally, we have compiled and collected TCR structures from the PDB (https://www.rcsb.org/) database, categorized based on binding and unbinding. Furthermore, there are TCR structures generated using TCRmodel2, and these all structures have undergone preprocessing to extract the CDR3β portion.

## Code Availability

The source code and model weights for TCRAD are available on GitHub (https://anonymous.4open.science/r/TCRAD-31E3) and on Zenodo (https://zenodo.org/records/10715856).

## Acknowledgments

This work was supported by National Natural Science Foundation of China grants (92374206 to P.Y., T2394512 and 12172371 to Y.Z.; 32090044 to J.L.), and Strategic Priority Research Program of the Chinese Academy of Sciences grant (XDB1000103 to J.L.). We acknowledge the Harbin Supercomputer Center and HPC-Service Station at the Center for Biological Imaging of the Institute of Biophysics for providing the computational resources.

## Contributions

J.L. conceived the project. C.L. and Y.G. Y. Z. designed the overall experimental and computational strategies. C.L. and Y.G. perform the data collection, wrote the Python scripts, built the AI-models and analyzed data. X. H. and Y. Z. purified proteins, constructed the cell lines, and performed T-cell function experiments. C. L. and Y. Z. prepared initial figures and wrote the first version of the manuscript. C. L., Y. G., Y.Z. and J. L. participated in the discussions and optimized figures. All authors contributed to revising the manuscript.

