## Supporting Methods for "TCRAD: An End-to-End Framework for Antigen-Targeted T Cell Receptor Design"

**Data Construction and Processing**

We initially retrieved sequence-level Peptide-CDR3β binding data from two databases: Tpp^1^ and Tchard^2^. Both datasets compile content from McPAS-TCR^3^, VDJdb^4^, and IEDB^5^ databases. In additon, Tpp incorporates data from 10ⅹ Genomics^6^, ImmuneCODE^7^, and TBAdb^8^, while Tchard additionally includes data from the MIRA^9^ set within the NetTCR-2.0 repository. Because both Tpp and Tchard augment negative samples through random peptide-TCR pairing, we retained only positive samples from Tpp, and retained both positive samples and experimentally validated negative samples from Tchard.

The collected data were filtered based on both the length distribution of the raw corpus and the model’s capacity constraints. Specifically, we retained peptides of length 8–15 amino acids and CDR3β sequences of length ≤20 amino acids, removed duplicates, and excluded clearly noisy or erroneous entries, including sequences containing non-standard amino acid characters, abnormal lengths, mismatched annotations, misaligned characters, or blank fields. After filtering, we obtained 157k positive samples and 300k negative samples. The 157k positive samples (denoted as K1-positive) were split into training, validation, and test sets at an 8:1:1 ratio for training the generation model. Fig. S1 shows the length distribution of the K1 dataset.

For data augmentation, we randomly paired 157 k positive samples to generate new negative samples. After removing duplicates among the random negatives, we obtained a peptide-CDR3β dataset termed K1, containing 519K entries. K1 was divided into training, test, and validation sets using the same 8:1:1 ratio to support robust model training and evaluation.

Additionally, we collected a binding dataset K3 from 10x Genomic^6^ to access zero-shot performance across different models. K3 contains complete new peptide-MHC-CDR3α-CDR3β pairs and is suitable for models with varying input requirements. After applying filtering criteria consistent with those used for K1, we randomly sampled 2,500 positive and 2,500 negative samples to create the K3 evaluation set. Overlapping samples between K1 and K3 were removed to ensure strict dataset independence. The final K3 dataset is entirely separate from K1 and share no common samples.

To train the Naturalness Classifier, we collected 600 k natural CDR3β sequences from the TABR^10^ dataset. Using randomly selected peptide as input for Sequence Generation (SG) module, we generated additional TCR sequences to expand the generated TCR dataset. This process continued until achieving a 1:1 ratio between natural to generated CD3β sequences. The resulting dataset, K2, was also divided into training, test, and validation sets using an 8:1:1 ratio. Optimal thresholds for both the Binding Classifier and the Naturalness Classifier were determined using ROC curve performance metrics on the K2 test set.

$${t^{*}=arg max}_{t}(Sensitivity(t)+Specificity(t)-1)$$

Where *t* denotes the decision threshold on the classifier score, $Sensitivity(t)$ is the true positive rate (TPR) at threshold *t*, and $Specificity(t)$ is the true negative rate (TNR). Equivalently, the false positive rate is $FPR(t)=1-Specificity(t)$. The optimal threshold $t^{*}$ was subsequently used to filter generated CDR3β sequences.

For the Structure Prediction module, we collected 260 bound TCR structures and 86 unbound structures from the PDB^11^ database. Corresponding CDR3β regions were extracted based on annotations provided by STCRDab^12^. Additionally, we utilized the TCRModel2^13^ tool to generate 5,834 TCR (derived from 10x Genomics^6^) structures to augment structural training. The Structure Prediction module was first pre-trained on generated structures and was then refined on experimentally resolved structures from the PDB. To ensure low homology, we performed sequence-based de-duplication on CDR3β and constructed the 8:2 train/test split such that no test-set CDR3β sequence had high sequence identity to any training-set sequence.

**Sequence Generation Module**

Inspired by the UniLM^14^ model, we adopted the methodologies and designed corresponding attention-mask patterns to train the Sequence Generation module. The UniLM model utilizes the classical Bert model as foundational architecture and introduces modifications to the mask matrix. A peptide of length M can be uniquely represented by a sequence of amino acid types denoted by p = ($p_{1}$, $p_{2}$, …, $p_{M}$). Similarly, a CDR3β of length L can be uniquely represented by a sequence of amino acid types denoted by c = ($c_{1}$, $c_{2}$, …, $c_{N}$). We developed an autoregressive protein language model capable of generating CDR3β sequences according to six user-defined generation orders:

$P(X)=\prod_{X \in unpredicted} P(X | c \in predicted ,pep)$	(2)

here X denotes the CDR3β position selected for prediction. The model predicts the amino acid at position X conditioned on the given peptide and the residues at all previously predicted CDR3β positions.

We leveraged the classical embedding strategy used in BERT, in which each sequence is represented through token embedding, positional embedding, and region embedding:

$$X_{embedding}^{0} = X_{token}^{0} + X_{position}^{0} + X_{region}^{0}$$

and

15-L times

$X_{token}^{0} =(E_{token}(p_{1})$,...,$E_{token}(p_{M})$,$E_{token}(PAD)$,$E_{token}(EOS)$,

$E_{token}(PAD)$,$E_{token}(c_{1})$,...,$E_{token}(c_{N})$,$E_{token}(PAD)$,$E_{token}(EOS))$

(20-N)/2 times

(20-N)/2 times

$X_{position}^{0} =(E_{position}(0)$,$E_{position}(1)$,...,$E_{position}(35)$,$E_{position}(36))$

$X_{region}^{0} =(E_{region}(0)$,...,$E_{region}($0$)$,$E_{region}(1)$,...,$E_{region}(1))$

Where $E_{token}$,$E_{position}$,$E_{region}$ are linear mapping functions correspond to embedding of residue types, residue positions, and sequence types. $X_{token}^{0}$,$X_{position}^{0}$,$X_{region}^{0}$ represent the resulting embedded vectors.

For position embedding, a total of 37 positions are ordered from 0 to 36. The first 16 positions represent peptide and an end-of-sequence (*EOS)* token, while the subsequent 21 positions represent CDR3β and *EOS* positions.

For region embedding, 0 represents peptide, and 1 represents CDR3β region. *PAD* and *EOS* signify sequence padding and sequence end tokens. In contrast to the traditional approach of padding sequences at the end, when padding the CDR3β segment of the sequence, we evenly distribute padding positions to both the beginning and end of the sequence. This strategy aims to align the central positions of higher importance in CDR3β, providing a distinctive approach to sequence padding.

After embedding, the processed sequence is passed into the Generation model, which selects the appropriate mask matrix based on the designated generation order. The mask matrix follows the principles below:

1. Each token is always allowed to attend to itself.
2. Peptide tokens form a conditioning context and may attend within the peptide segment, but cannot attend to any CDR3β tokens.
3. When predicting a CDR3β position, the token may attend to all peptide tokens and to previously generated CDR3β positions, but cannot attend to any not-yet-generated CDR3β positions. For mutation prediction, CDR3β tokens may additionally attend to non-mutated CDR3β positions as known context.
4. PAD and EOS tokens are masked from CDR3β positions and are restricted to interact only within the peptide segment. This prevents information leakage through special tokens from not-yet-generated CDR3β positions.

Following these principles, we constructed different mask attention matrices according to the generation order. The attention matrix for CDR3β form mutation-design tasks is shown in Supplementary Fig. S2.

After the generated sequence undergo embedding and mask application through the SG network, the model outputs the probability distribution over candidate amino acids at the target position. Here, we employ a hybrid decoding strategy that combines nucleus (top-p) sampling and top-k sampling^15^. Initially, the decoded MASK predictions are sorted in descending order. Utilizing the TOPP strategy, candidates are selected whose cumulative probability reaches 80% of the total sum. Subsequently, the TOPK strategy is applied to further filter these candidates. Specifically, for different positions, considering the varying importance of CDR3β positions in peptide-CDR3β binding tasks based on previously analysed attention weights, different conditions are applied to determine the number of retained candidate items. Ultimately, if the filtered candidate set is empty, the top two items from the original candidate set are used. This hybrid strategy dynamically adjusts the generated candidate set based on specific circumstances, catering to diverse requirements. Pseudocode is provided in Supplementary Algorithm 1.

**Model Training Details**

We initiated the training of the Binding Classifier model using the K1 dataset, concurrently engaging in pre-training tasks involving both Masked Language Model (MLM) and Binding Prediction. During pre-training, we randomly masked three peptide-CDR3β positions, including *PAD* and *EOS* positions, for predicting masked positions and binding reactivity. Subsequently, the pre-trained model was utilized for two main objectives. Firstly, solely focusing on Binding Prediction (with zero masked positions) resulted in the development of the Binding Classifier, facilitating the analysis of the importance distribution of CDR3β positions. Secondly, using the pre-trained model as a foundation, MLM task training on the K1-positive set was conducted to obtain the Generation model. Drawing inspiration from the training strategy of the UniLM model, we have enhanced the training policy for addressing the CDR3β generation problem (Algorithm 1).

For each epoch, we employed six distinct sequence generation strategies as mentioned in Fig 3, allocating one-sixth of the training tasks to each model. The loss for each epoch was the sum of the model losses across the six training tasks. Regarding the masking strategy, we specifically masked three positions (including the *EOS* and *PAD* positions) in the CDR3β region. In 80% of cases, direct masking with MASK was applied, in 10% of cases, a random residue was chosen to replace the mask, and in the final 10%, the real residue was retained without replacement.

In addition, the 80% case of masking involved independently masking the three positions, while the remaining 10% involved either masking two consecutive positions (bigram) and one independent position or masking three consecutive positions (trigram). For the Naturalness Classifier, we initially performed MLM tasks on individual CDR3β sequences, followed by prediction tasks for Natural Prediction. Cross-entropy served as the loss function for MLM tasks, while Mean Squared Error (MSE) was employed for Prediction tasks:

$$Binding Prediction Loss = \frac{1}{n}\sum_{i=1}^{n} {(y_{ture,i}-y_{pred,i})}^{2}$$

Where $n$ represents the total number of samples. $y_{ture,i}$ is the ground truth value for the $i$*-th* sample, $y_{pred,i}$ is the predicted value by the model for the $i$*-th* sample.

$$Residue Classfication Loss = -\sum_{i} y_{ture,i}\times\log(y_{pred,i})$$

Where $i$ is the index representing the residue type,$y_{ture,i}$represents the actual probability of belonging to the $i$*-th* residue type. $y_{pred,i}$ is the model's predicted probability for the $i$*-th* residue type.

Regarding the Structure Module, due to the relatively short length of the CDR3β sequence, the complexity of structure prediction is comparatively lower. To adapt to this situation, we reduced the number of network layers and decreased the embedding dimensions. The training of the Evoformer layer and Structure model layer was conducted based on the previously trained Naturalness Classifier and Binding Classifier. We performed joint training of the two classifiers to train the overall model. Initially, the model underwent pre-training on TCRModel2-generated Unbounded TCR structures. Subsequently, fine-tuning was carried out on natural structure data. Specifically, within an epoch, we randomly selected Bounded and Unbounded structures for prediction tasks, with both tasks sharing the weights of the Structure model and Evoformer. Subsequently, model fine-tuning was conducted separately for each task. In addition, we refined the AlphaFold2^16^ structural violations loss to better capture local peptide-bond geometry. Specifically, we introduced additional loss terms on bond lengths and bond angles involving the CA, C, O, and N atoms, thereby enforcing peptide-bond planarity. In the original AlphaFold2 formulation, the carbonyl oxygen is not explicitly constrained in the structural violation loss, which can lead to non-coplanar backbone geometries that violate the planar nature of the C=O double bond—an issue that is particularly pronounced for short, flexible CDR3β loops.

**In Vitro Experimental TCR Library Construction**

First, we used the sequence from PDB entry 2BNR, corresponding to 1G4 TCR, as the backbone for constructing the experimental TCR library. Using the corresponding NY-ESO-1 sequence SLLMWITQC as a input for our model, we conducted full design and mutation optimization for the CDR3β sequences. During mutation optimization, mutation sites were randomly sampled within the CDR3β, and the Sequence Generation Module was then used to generate a diverse set of candidate CDR3β sequences.

For all generated sequences, we initially applied the Naturalness Classifier to remove sequences with low naturalness scores. Subsequently, the Binding Classifier was used to further exclude sequences unlikely to bind the pMHC complex. We also referenced the TABR-BERT^10^ and TEIM^17^ models to exclude sequences with excessively low binding potential. For the remaining sequences, we conducted further filtering based on sequence similarity and the technical feasibility of sequence synthesis. Ultimately, the resulting candidate pool was down-selected to a high-confidence library of 29 designed TCRs, presented in Supplementary Table S5.

**TCR Binding and Functional Assay**

***Cell lines and Culture***

293FT cells and J76 cells were used in this study. The 293FT cell line, used for lentivirus packing, were cultured in DMEM (MeilunBio) supplemented with 10% fetal bovine serum (MeilunBio), 1% penicillin-streptomycin (Gibco) and 0.01% plasmocin^TM^ treatment (InvivoGen). The J76 cells, which lack endogenous TCR expression and were used to express the engineered TCRs, were cultured in RPMI 1640 (MeilunBio) supplemented with 10% fetal bovine serum (MeilunBio), 1% penicillin-streptomycin, 1mM sodium pyruvate ([Sigma-Aldrich](http://www.labgogo.com/BrandInfo/BrandShop.aspx?pn=sigma-aldrich)), and 0.01% plasmocin^TM^ treatment.

***Lentivirus Production and Transduction***

First, the designed CDR3β sequences were cloned into the pHAGE lentiviral vector, which contains the 1G4 TCR and a BFP reporter linked via an IRES sequence. The resulting plasmid was co-transfected into 293FT cells together with the packaging plasmids psPAX_2_ and pMD2.G, using the jetPRIME® transfection reagent (Polyplus). After 48 hours of incubation, the viral supernatant was harvested, centrifuged to remove cell debris, and the clarified supernatant was used to transduce J76 cells. The culture medium was replaced after one day, and transduced cells were enriched by flow cytometry sorting three days post-infection.

***Peptides and Tetramers***

Biotinylated HLA-A2/NY-ESO-1 was purified as described in a previous work^18^, and stored at -80°C. Streptavidin (SA) labeled Alexa Fluor 647 (Thermo Fisher Scientific) was mixed with biotinylated HLA-A2/NY-ESO-1 at 4°C to form Alexa Fluor 647-labeled pHLA tetramers.

***NFAT-GFP Reporter Assay***

J76 cells were transfected with an NFAT-GFP reporter plasmid prior to lentiviral transduction of the TCR constructs. A total of 2×10^4^ TCR-expressing J76 cells were seeded into wells pre-coated with immobilized pMHC complexes. After 24 hours of stimulation, NFAT activation was quantified by flow cytometry. J76 cells lacking TCR were used as the negative control, and that transfected with 1G4 TCR were used as the positive control.

***Flow Cytometry***

Successfully infected J76 cells were sorted based on BFP reporter expression using a BD Aria cell sorter. TCR surface expression was assessed using a fluorophore-conjugated antibody hCD3ε-PE (Invitrogen, #2939147). Alexa Fluor 647-labeled pHLA tetramers were utilized to evaluate the binding of the expressed TCRs to their cognate pHLA complexes. The percentage of GFP-positive cells in the TCR-expressing J76 cells co-cultured with pHLA tetramers was quantified by flow cytometry as a measure of T cell activation. All flow cytometry data were analysed using FlowJo software.

**Details of Baselines**

Below, we provide additional details on how each baseline model was adapted and utilized for comparison. All default hyperparameters were retained as defined in the original implementations. For each method, our dataset was reformatted to match the required input specifications. Because different TCR-antigen binding prediction models accept different combinations of input sequences, TCRAD, Panpep, and TEIM^17^ use peptide and CDR3β; pMTnet additionally requires MHC type; and TABR further incorporates CDR3α — we curated the K3 dataset to include only samples with complete peptide-MHC-CDR3α-CDR3β information, ensuring compatibility across all baseline models. Pre-trained model weights provided by the original implementations were used without modification.

Both **TEIM**^17^ and **TABR-Bert**^10^ output binding scores ranging from 0 and 1. In contrast, **pMTnet**^19^ does not directly output a binding probability, instead, it reports a percentile rank that reflects the relative binding strength of the given TCR-pMHC pair compared to 10,000 randomly sampled TCRs against the same pMHC. To ensure consistency with other baselines, we modified the original code to extract the underlying predicted score as the model’s output. **Panpep**^20^ includes few-shot learning, zero-shot learning, and majority learning frameworks. To maintain a fair comparison under a zero-shot evaluation protocol, we used the zero-shot mode for Panpep’s predictions.

**Implementation Details**

***Framework and Hardware***

Our models were implemented using PyTorch and trained using the Adam optimizer with a learning rate of 1 × 10^−4^. To facilitate stable convergence, we employed a dynamic learning rate schedule that reduced the learning rate by a factor of 0.1 whenever the validation loss failed to improve for one epoch. All experiments were conducted on eight NVIDIA V100 GPUs. To avoid overfitting, we employed an early stopping strategy with a patience of 4 epochs, based on validation loss monitoring.

For the Binding Classifier and the Generation Model, the batch size was set to 256, whereas the Naturalness Classifier was trained with a batch size of 512. All models employed an 8-layer, 6-head Transformer encoder architecture. For structure prediction, we use an 8-layer, 12-head Structure Module together with an 8-layer Evoformer. Due to the limited size of the structure dataset, we utilized a mini-batch learning strategy, updating the model parameters after every four samples.

***Decoding strategy of Generation Module***

Algorithm 2 describes the procedure used to decode and determine the final candidate amino acid at each position. After the generation module outputs a probability distribution over all amino acid types for the target position, the decoding strategy selects the final residue based on a combination of probability filtering and position-specific constraints. This process ensures that the chosen amino acid is both statistically plausible and consistent with the positional requirements of CDR3β sequence design.

**Algorithm 1:** Training algorithm of Generation Module

**Input**: Training dataset K1-Positive-train, validation dataset K1-Positive-val

**Output**: Trained model parameters

1： Load training dataset and validation dataset
2： Initialize BERT model with pretrained parameters θ_0
3： Set optimizer and loss function
4： for epoch e = 1 to N do
5： for each batch (X, y) in K1-Positive-train do
6： Generate masked input and selecting random mask strategy
7： Compute model predictions y_hat
8： Compute masked language modeling loss L_MLM
9： Perform backpropagation and update theta using gradient descent
10： end for
11： Compute validation loss L_val on K1-Positive-val
12： Adjust learning rate using scheduler
13： if validation loss increases for 3 consecutive epochs then
14： break
15： end if
16： end for
17： Saving the trained model

**Algorithm 2: Hybrid Decoding Strategy for Candidate Selection**

**Input**: Predicted scores over amino acid types (Arr_ori) and decoding position (Pos)

**Output**: Candidate amino acid probabilities (C_prob) and types (C_type)

1： Set temperature to 1.
2： Rank amino acid probability Arr_Rank=sorted(softmax(Arr_ori) / temperature)
3： Perform top-p selection with threshold 0.8
4： Remove C_prob with probability < 0.05
5： Perform top-k selection:
6： if Pos in [The most critical sites]:
7： Keep up to 6 candidates, C_prob[:6]
8： elif Pos in [Secondary important sites]:
9： Keep up to 3 candidates, C_prob[:3]
10： else:
11： Remove C_prob with probability < 0.5 * max(C_prob)
12： Keep up to 2 candidates, C_prob[:2]
13： if C_prob empty:
14： C_prob = Arr_Rank[:2]
15： Derive the corresponding amino acid types C_type from C_prob.
16： return C_prob, C_type

Note: The most critical sites and secondary important sites are defined based on the weights in Fig. 5, and their relative sizes can be adjusted accordingly.
