## Supporting Figures for "TCRAD: An End-to-End Framework for Antigen-Targeted T Cell Receptor Design"

**Supporting Figures and Legends**

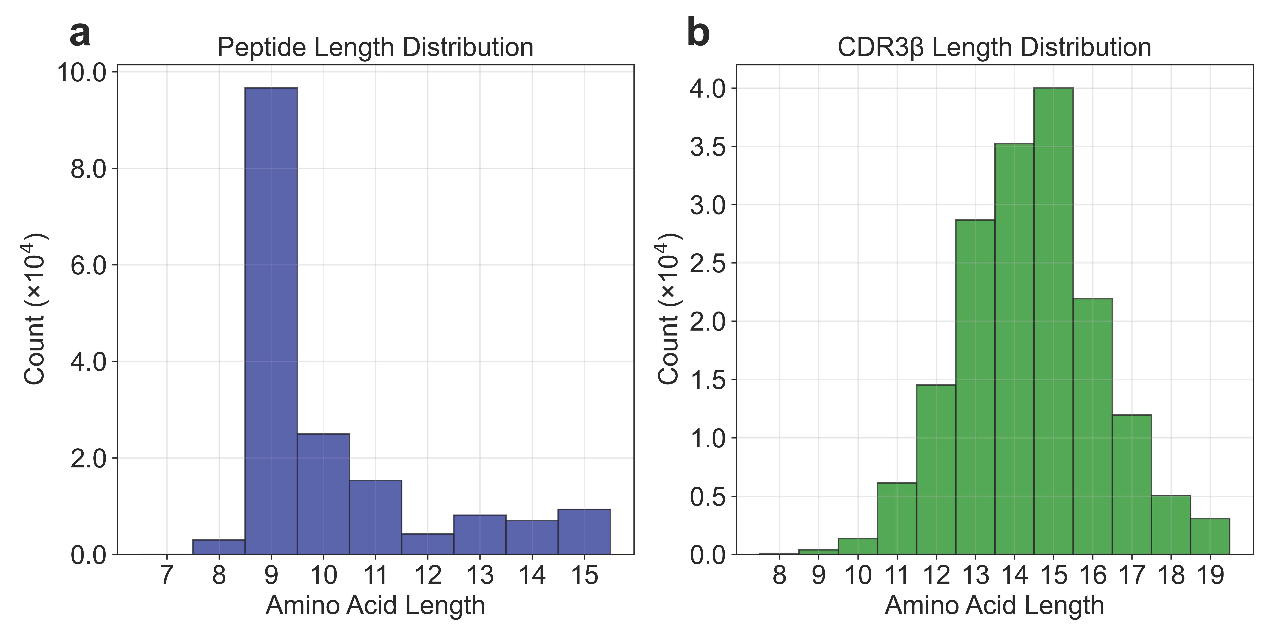

**Fig. S1: Length distribution of K1.** The figure presents the length distribution of Peptide and CDR3β sequences in the K1 dataset after sequence filtering and noise removal.

**a**. CDR3β Length Distribution: The distribution shows a peak around length 12, with most sequences falling between 10 and 16 amino acids.

**b**. Peptide Length Distribution: The distribution reveals that the majority of peptide sequences are between 8 and 10 amino acids, with a concentration at length 9.

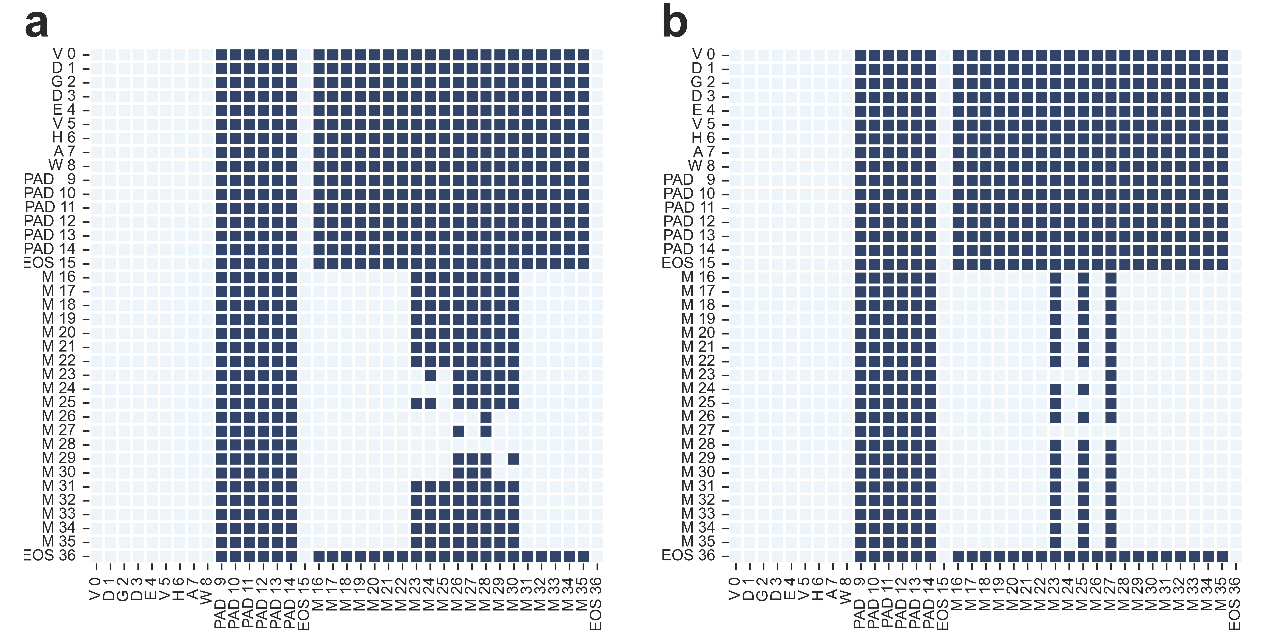

**Fig. S2: Mask of Mutation for Optimization Design.** Building upon the MASK strategy derived from the importance of CDR3β positions.

**a**. The generation prediction is performed for the contiguous middle 8 residues of the CDR3β sequence, given the two end segments of CDR3β. e.g. ------XXXXXXXX------ (given the length of CDR3β), ---CASSXXXXXXXXFF--- (given the starting and ending residues of CDR3β).

**b**. Mutations are applied to the CDR3β residues to achieve optimization, e.g. ----CASXSXGXQYF-----(optimizing three residues).

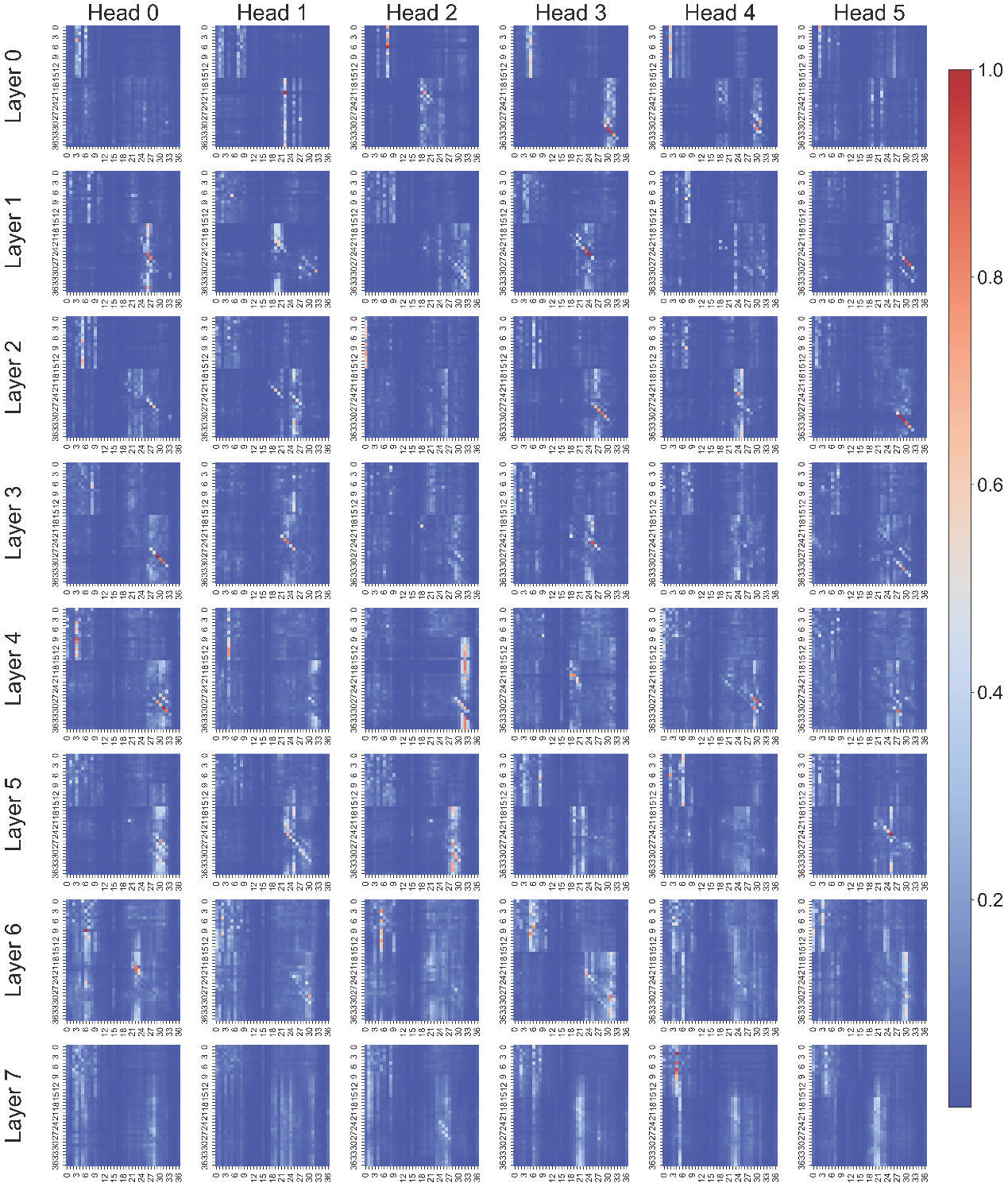

**Fig. S3. Analyzing the weights on the K1-Test, examining the positional weight information for the P-CDR3β Binding task. We visualize the learned positional weight distribution of the binding discriminator when evaluating Peptide-CDR3β pairs in K1-Test, highlighting how different peptide and CDR3β positions contribute to the binding score. The analysis quantifies position-dependent importance captured by the model during inference, consistent with the binding module design.**

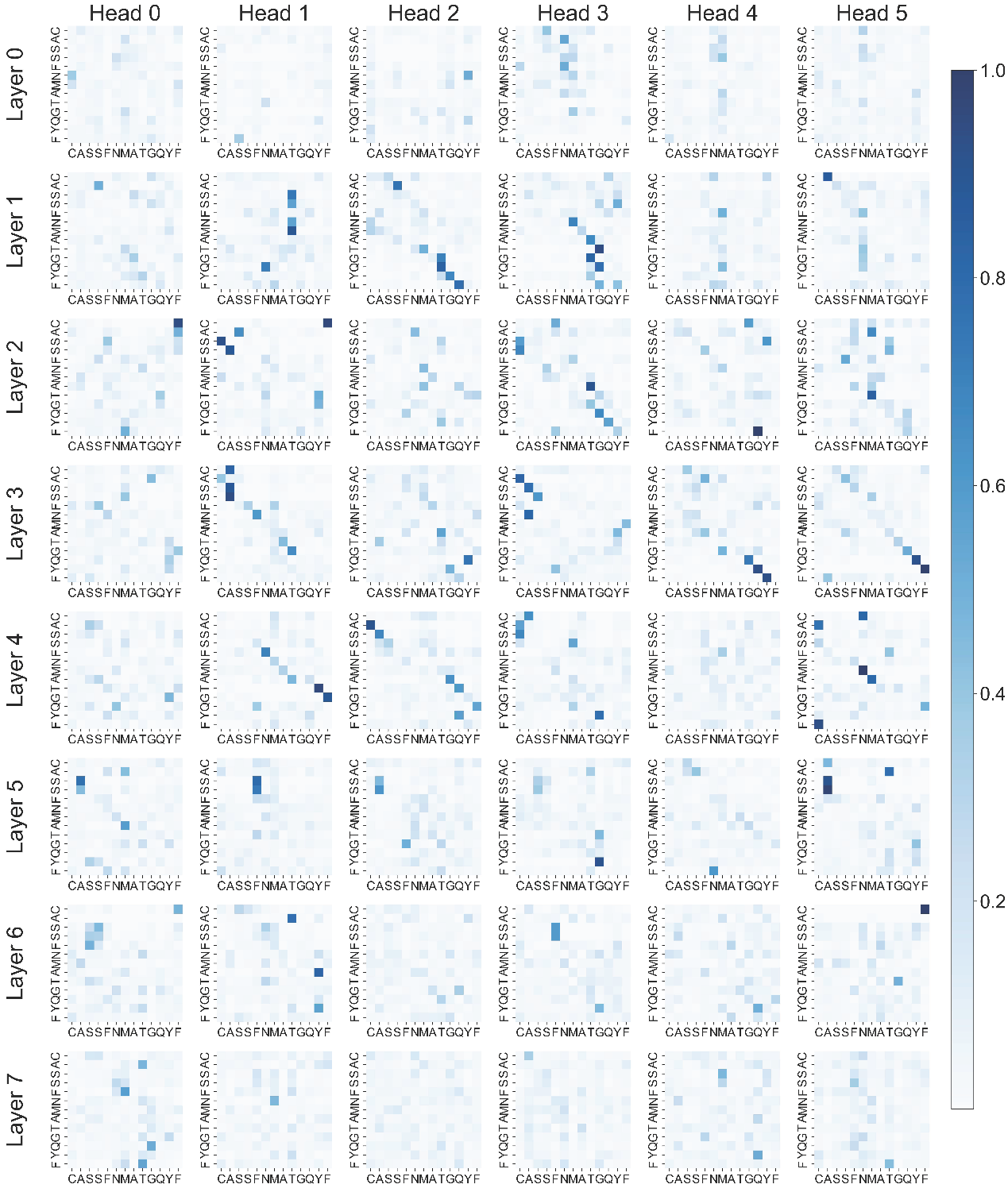

**Fig. S4. Inputting the CDR3β (CASSFNMATGQYF) corresponding to PDB: 5bs0 and the Peptide (ESDPIVAQY) into the Binding Classifier, obtaining the attention weight matrix of the CDR3β portion, with the model being an eight-layer six-head architecture, extracting the CDR3β portion from all self-attention layers.**

### Table S1 Dataset statistics of training datasets.

| **DataSet** | **Positive** | **Negative** | **Total** | **Source** |
| --- | --- | --- | --- | --- |
| K1 | 157K | 362K | 519K | Tpp^1^  Tchard^2^ |
| K2 | 600K | 600K | 1.2M | Self-Generated  TABR^3^ |
| K3 | 2.5K | 2.5K | 5K | 10x Genomics^4^ |
| Natural TCR Structure | NA | | 260-Bounded  86-Unbounded | PDB^5^  STCRDab^6^ |
| Predicted TCR Structure | NA | | 5834 | TCRModel2^7^ |

**Table S2: Optimization of the middle TRAGDT position based on CASSTRAGDTQYF for the peptide GILGFVFTL.**

The "clsf" column represents the prediction values from the Binding Classifier, while the "dis" column represents the prediction values from the Naturalness Classifier. Both values range from 0 to 1. A higher value in the "clsf" column indicates a higher likelihood of binding, whereas a lower value in the "dis" column suggests better naturalness.

The table is provided as a separate file: GILGFVFTL_CASSTRAGDTQYF.csv

**Table S3: Mutation optimization based on CASSQSPGGTQYF for the peptide GLCTLVAML, with optimization of the middle QAGGGI position.**

The table is provided as a separate file: GLCTLVAML_CASSQSPGGTQYF.csv

**Table S4: Designing with a complete regeneration approach for the NY-ESO-1 antigen.**

The table is provided as a separate file: Full_Design_for_2BNR_SLLMWITQC.csv

**Table S5: TCR library constructed for biological experimental validation.**

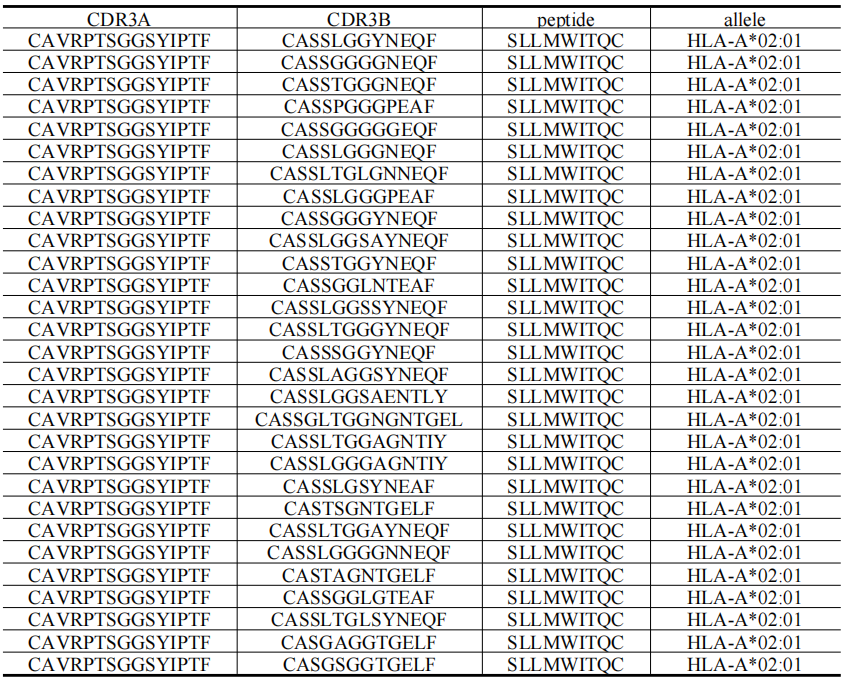

The table is also provided as a separate file: library.xlsx
